# Integration of Xeno-Free Single-cell Cloning in CRISPR-mediated DNA Editing of Human iPSCs Improves Homogeneity and Methodological Efficiency of Cellular Disease Modelling

**DOI:** 10.1101/2022.04.12.488094

**Authors:** Atefeh Namipashaki, Kealan Pugsley, Xiaodong Liu, Kirra Abrehart, Sue Mei Lim, Guizhi Sun, Marco J. Herold, Jose M. Polo, Mark A. Bellgrove, Ziarih Hawi

**Affiliations:** Turner Institute for Brain and Mental Health, School of Psychological Sciences, Monash University, VIC, Australia; Department of Anatomy & Developmental Biology, Monash University, Melbourne VIC 3800, Australia. Development and Stem Cells Program, Monash Biomedicine Discovery Institute, Melbourne VIC 3800, Australia. Australian Regenerative Medicine Institute, Monash University, Melbourne VIC 3800, Australia; Walter and Eliza Hall Institute of Medical Research, Melbourne, VIC, Australia. Department of Medical Biology, University of Melbourne, Melbourne, VIC, Australia; Adelaide Centre for Epigenetics and the South Australian Immunogenomics Cancer Institute, The University of Adelaide, SA, Australia

## Abstract

The capability to generate induced pluripotent stem cell (iPSC) lines, in tandem with CRISPR-Cas9 DNA editing, offers great promise to understand the underlying genetic mechanisms of human disease. The low efficiency of available methods for homogeneous expansion of singularised CRISPR-transfected iPSCs necessitates the coculture of transfected cells in mixed populations and/or on feeder layers. Consequently, edited cells must be purified using labour-intensive screening and selection, culminating in inefficient editing. Here, we provide a xeno-free method for single-cell cloning of CRISPRed iPSCs achieving a clonal survival of up to 70% within 7-10 days. This was accomplished through improved viability of the transfected cells, paralleled with provision of enriched environment for the robust establishment and proliferation of singularised iPSC clones. Enhanced cell survival was accompanied by a high transfection efficiency exceeding 97%, and editing efficiencies of 50-65% for NHEJ and 10% for HDR, indicative of the method’s utility in stem cell disease modelling.

## INTRODUCTION

Despite advances in our ability to detect DNA variants conferring risk to complex genetic conditions, our knowledge of their functional effects on biological systems is lagging (Gallagher and Chen-Plotkin, 2018). Commonly used disease models, such as animal and cancerous cell lines, do not faithfully resemble disease-relevant genotypic and phenotypic properties (Hoffmann et al., 2020). Furthermore, inaccessibility of certain living tissue (e.g., brain) necessitates the development of relevant experimental models to help understand the molecular mechanisms through which genetic variants influence disease states (Kampmann, 2020). Induced pluripotent stem cells (iPSCs) provide a valuable alternative to these methods. When obtained from patients, iPSCs are genetically enriched for the presenting disease, recapitulating the complex aetiology of the target condition (Matos et al., 2020). However, comparing patient-derived iPSCs to those derived from healthy controls is often impractical. Without large sample sizes, genetic heterogeneity limits the statistical power of case-control comparisons, making it difficult to correlate phenotypic differences to disease status (Matos *et al*., 2020). The combined use of iPSC and CRISPR-Cas9 DNA editing can assist to reduce this heterogeneity through generation of isogenic cell lines. Differing only by a genetic sequence(s) of interest, this approach significantly reduces the number of samples required to identify the effect of disease-associated genetic variants (Chang et al., 2018; Soldner et al., 2011).

The CRISPR-Cas9 machinery mediates genome editing by inducing a double strand break (DSB) in a specified genomic locus, resulting in activation of either the non-homologous end joining (NHEJ) or homology-directed repair (HDR) cellular repair pathway (Sander and Joung, 2014). While the error-prone NHEJ repair system can introduce indels at the cut site, a precise HDR event can be induced through the provision of a template DNA sequence. Increasing the probability of an intended on-target edit has been an active area of research in recent years, with specific focus on improving the guide RNA (gRNA) and template donor molecule design, as well as development of strategies to enrich for CRISPRed cells with the correct edit (Ikeda et al., 2018; Kwart et al., 2017; Merkle et al., 2015; Mianne et al., 2020; Steyer et al., 2018). Nonetheless, at present, CRISPR-mediated editing frequently results in variable catalytic events, producing a genetically heterogeneous population of edited cells following delivery of the CRISPR-Cas9 complex (Simkin et al., 2022). Clonal expansion of individual CRISPRed cells is therefore necessary to produce a cell line containing only the desired on-target edit. This is usually achieved through limiting dilution or single-cell culturing following assortment of individual cells via fluorescence-activated cell sorting (FACS). The resultant monoclonality increases the odds of capturing the intended on-target edit and ensures homogeneity of the edited cells. This restricts the source of phenotypic differences between isogenic cell lines to the target mutation (Mianne *et al*., 2020), permitting valid conclusions as to the consequences of the intended genetic determinant on processes of biological functioning.

One of the major bottlenecks hampering the application of CRISPR-Cas9 in iPSCs is homogenous single-cell cloning following editing. Unlike non-primary cell types such as HEK-293 and SH-SY5Y, culturing CRISPRed iPSCs from a single-cell state into a stable clone is a challenging task. This is consequence of the innate susceptibility of iPSCs to dissociation-induced apoptosis and elevated risk of p53-mediated apoptosis in response to the toxicity of the CRISPR-Cas9 complex (Das et al., 2020; Ihry et al., 2018; Ohgushi et al., 2010). To increase survival, transfected cells are therefore cultured and expanded in mixed populations, commonly with a low density that can still assure viability. This results in the generation of mosaic colonies, requiring purification by either antibiotic selection or successive rounds of screening (e.g., droplet digital PCR, T7EN1 assay, Surveyor assay) and sorting to increase the percentage of cells with the desired edit within a given population (Li et al., 2016; Miyaoka et al., 2014; Yumlu et al., 2017). Though these approaches help to enrich for the target mutation, the purified cells are the progeny of mixed colonies and as such possess incomplete isogeneity, complicating subsequent phenotyping. Alternate methods may circumvent this heterogeneity through application of feeder layers (e.g., mouse embryonic fibroblasts) to support the growth of successfully edited single-cells. However, these cannot be considered xenogeneic free (Byrne et al., 2014; Singh, 2019), and are suboptimal for modelling human genetic disease mechanisms.

Here, we describe the development of a novel method for feeder-free establishment and expansion of singularised CRISPRed iPSC clones homogeneous for targeted DNA edit(s). This has been achieved through improvement of the viability of transfected cells following FACS and provision of an enriched environment for the robust proliferation of singularised iPS cells. Practical application of our proposed method in three cell lines derived from various parental cell types achieved up to 70% single-cell clone survival post-transfection, accompanied by >97% transfection efficiency. Editing was observed across 50 to 65% of surviving clones in NHEJ and 10% for HDR.

## RESULTS

### Utilizing ribonucleoproteins (RNPs) and lipid delivery can increase iPS cell survival post-transfection: a prerequisite to improving single-cell clonal expansion

Successful single-cell cloning of CRISPRed iPSCs is primarily dependent on the viability of the transfected cells. Accordingly, we made a series of methodological considerations with a focus on increasing cell survival. The CRISPR experiments in this study were performed on three different iPSC lines generated in-house from fibroblast (HDFn and HDFa) and peripheral blood mononuclear cell (MICCNi002-A) lineages. The editing complexes were designed to target intron 6 of *HPRT* and exon 1 of *FOXP2* in the HDFn/HDFa and MICCNi002-A lines, respectively. To eliminate the impact of DSB, a non-targeting guide RNA control was also used to determine post transfection iPSC cell survival.

One of the first determinants of cell viability is vector selection, with DNA-based vectors such as plasmid delivery conferring high degrees of cellular toxicity. For this reason, we opted to use a protein-based system of ribonucleoproteins (RNPs) consisting of a gRNA and tracrRNA duplex and the Cas9 enzyme. Such approaches have been previously reported to result in up to two-fold more edited colonies than achieved via plasmid transfection (Kim et al., 2014). Toxicity associated with the delivery of the CRISPR complex into the intracellular environment represents another significant barrier to achieving high cell viability. Electroporation is a commonly used method which physically delivers the editing machinery into cellular compartments using electrical shock, with high transfection efficiency (Lino *et al*., 2018). This contrasts with chemical lipofection, which instead utilises a cationic lipid nanoparticle encapsulating the gRNA: Cas9 complex to effectively bypass the negatively-charged cell membrane. While most studies have focused on evaluating the transfection efficiency of these two techniques, their influence on post-transfection iPS cellular viability has been less frequently addressed (Kim and Eberwine, 2010; Sharifi Tabar et al., 2015; Zizzi et al., 2010). Hence, we performed a CRISPR experiment for the delivery of the non-targeting RNPs into the three different iPSC lines to compare cell survival between the methods. For electroporation, a mixture consisting of RNPs, the Nucleofector electroporation solution, and singularised iPSCs were added to the 4D Lonza Nucleofector system according to the manufacturer’s instructions. This nucleofector system is designed to direct the CRISPR complex into the cell nucleus and thus is more appropriate for hard to transfect cell lines such as iPSCs (Hamm et al., 2002). Using the lipofection method, the RNPs were mixed with the commercially available lipofectamine stem transfection reagent and subsequently combined with iPSCs dissociated into small clumps of 3-10 cells. Post transfection cell survival was assessed using FACS. Figure 1A depicts the cell survival rate for the three iPSC lines. Though the average viability for the electroporation was measured at 46.84% ±1.18, the lipofection method demonstrated at least 40 percent higher cell survival (*p* = 0.001), with an average cell survival rate of 86.14% ± 0.51. The lipidic delivery system was therefore applied in subsequent CRISPR experiments.

**Figure 1.**
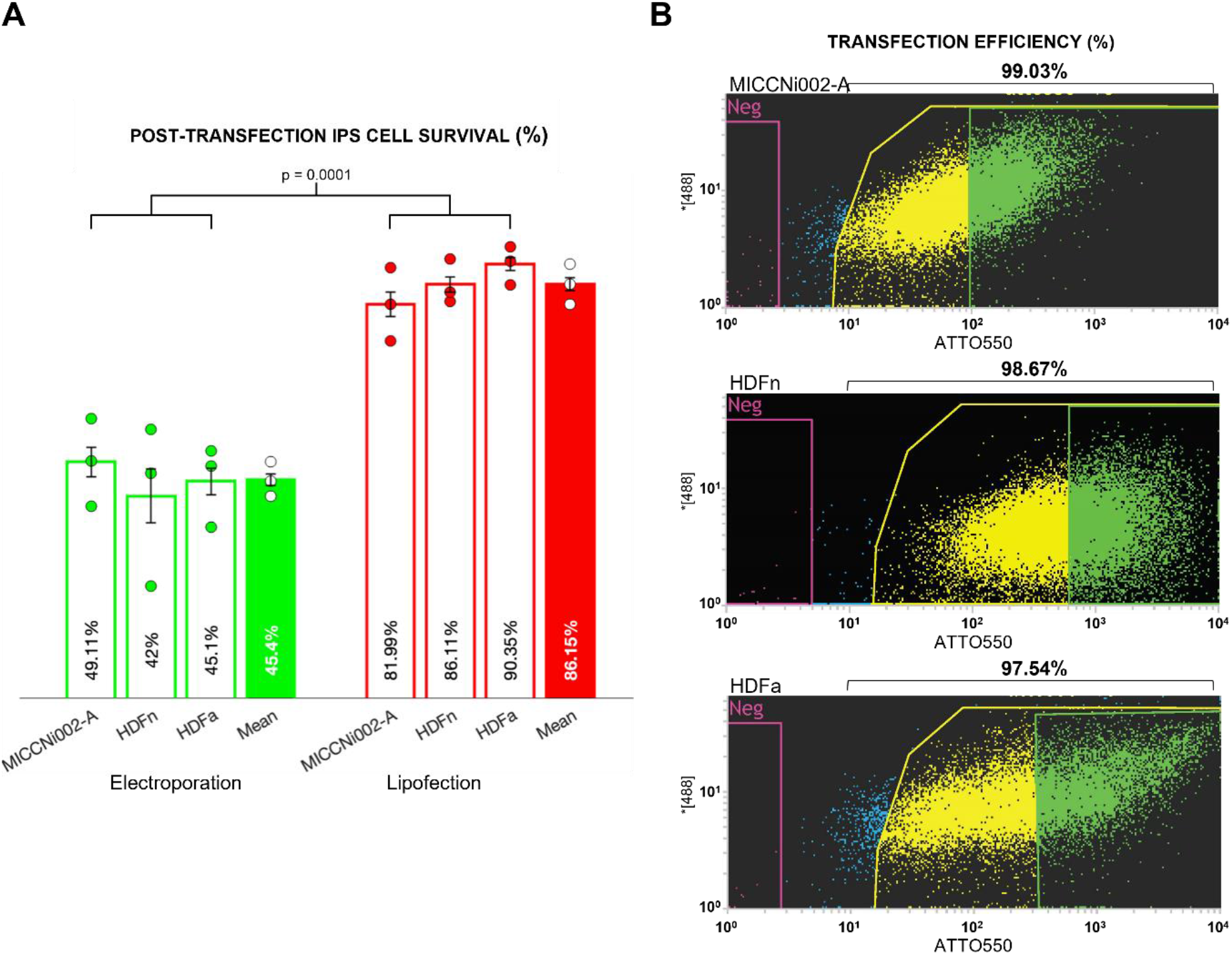
RNP lipid delivery achieves high cell survival and transfection rate. **(A)** Flow cytometry analysis of CRISPRed iPSCs transfected using the lipid based CRISPR delivery system shows a substantially higher degree of cell viability relative to electroporated cells, exceeding 40% (p = 0.0001). **(B)** Flow cytometry dot plot demonstrates that lipid RNP transfection using Lipofectamine Stem Transfection Reagent achieves close to 100% transfection efficiency (yellow and green) across three independent iPSC lines. The ATTO 550 (X-axis) was plotted against a short wavelength laser channel (488; Y-axis) to achieving a clear separation of the negative and positive population. Un-transfected cells are shown in purple (< 3%). Reproducibility of the transfection efficiencies were tested across three independent experiments.

### Combined application of RNP vector selection and lipid-based delivery of the CRISPR machinery yielded high transfection efficiency in human iPSCs

Successful delivery of the DNA-editing machinery into the intracellular environment is a major factor affecting the efficacy of CRISPR-based editing (Yang et al., 2014). Logically, where high rates of successful transfection can be achieved in tandem with persistent survival of CRISPRed cells, the probability of obtaining the desired edit is significantly enhanced.

To confirm that the combined RNP-lipid delivery approach yields a high transfection efficiency alongside enhanced cell viability, we evaluated the RNPs conjugated with the ATTO-550 uptake via FACS. Lipofection of the RNP complexes resulted in a transfection efficiency exceeding 97% (**Figure 1B**) in the three iPSC lines, greatly improving upon those efficiencies previously reported using other delivery and vector systems (Geng et al., 2020; Singh, 2019; Wang et al., 2017; Xu et al., 2018; Yumlu *et al*., 2017).

### Our unique post-transfection single-cell cloning method demonstrates high clone survival while preserving genetic homogeneity

Isolation of the healthy viable CRISPR-transfected cells through FACS as 1 cell/well is the next step for the establishment of genetically homogeneous iPSC clones. However, culturing of singularised iPSCs can be difficult in the absence of supportive feeder layers. Three major factors influence iPS cell survival under feeder-free conditions: (1) the cell culture media, (2) utilisation of small molecule inhibitor supplements, and (3) the surface matrices (Li et al., 2009; Watanabe et al., 2007). As such, we performed a gradual optimisation approach focusing on these three elements, comparing different commercially available products to evaluate their combined influence on cell survival in the MICCNi002-A line targeting exon 1 of *FOXP2*. In the first iteration of the optimisation process, single transfected iPSCs were sorted in individual wells of a 96-well plate pre-coated with Vitronectin (VTN) containing Essential 8 Flex medium and supplemented with RevitaCell. After 10-15 days, we observed a survival rate of 12.5% ±0.01. Substitution of RevitaCell with CloneR supplement enhanced the frequency of emerging single-cell clones to 22.57% ±0.01, which was further increased to 36.46% ±0.006 with the additional substitution of E8 Flex medium with StemFlex medium. The highest degree of single-cell clone survival was observed when combining the application of StemFlex+CloneR with the human Laminin-521 surface matrix in place of VTN, resulting in 70.14% ±0.03 of positively CRISPR-transfected iPSC clones emerging within 7-10 days (**Figure 2A**). The time periods represented for each condition are the ranges between which the majority of single-cell clones for a given condition reached confluency for further analysis. Day 10 was selected as the optimal shared timepoint to count the cells in all conditions. Application of this method in each of the HDFn, HDFa, and MICCNi002-A lines resulted in a post-transfection survival frequency ranging between 61.8 to 70.1%, demonstrating consistently high-rates of survival between lines. To account for the effect of CRISPR-Cas9 toxicity on cell viability, FACS-sorted cells of the parental lines were subject to the StemFlex+CloneR+Laminin-521 plating conditions, achieving a 7-10 day clonal rate exceeding 86% (**Figure 2B**). This suggests that ∼20% of cell death observed in our CRISPR-positive cell populations is the direct consequence of the introduction of the editing machinery.

**Figure 2.**
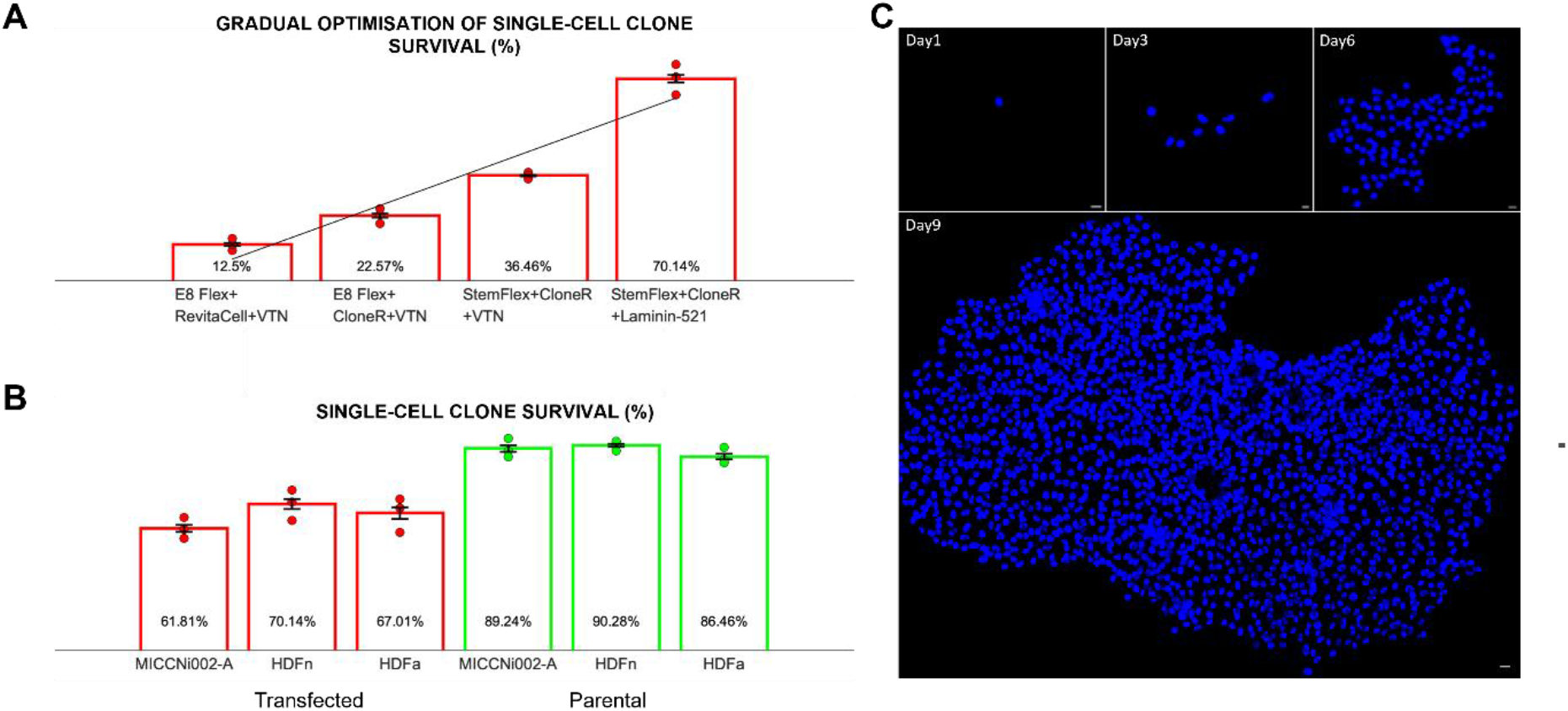
Integration of RNP lipid delivery with our unique method of single-cell cloning achieves highly improved homogenous CRISPRed-iPSC clone survival. **(A)** Gradual optimisation method shows utilisation of a rich environment containing StemFlex and CloneR supplement in wells coated with Laminin-521 achieves the highest single-cell clonal emergence following CRISPR transfection of up to 70.1%. **(B)** Application of the StemFlex+CloneR+Laminin-521 plating conditions in three different cell line led to consistent high-level survival in CRISPRed (61.8 to 70.1%) and non-CRISPRed single-cell clones (86.4 to 90.2%). (C) Microscopy images showing a single-cell clone growth over time. Cells were imaged via the fluorescent NucBlue Live cell stain.

Together, these findings demonstrate that the combination of these reagents can significantly improve the establishment of genetically homogeneous single-cell clones (**Figure 2C**), countering the degree of cell death conferred by the CRISPR complex to achieve high rates of cell survival.

### The frequency of genome editing following application of our method shows high consistency after single-cell cloning

Double strand breaks caused by the Cas9 enzyme can induce p53 cascade mediated apoptosis, which, together with decreased cellular viability in singularised transfected iPSCs, may compound cell susceptibility to death following single-cell isolation (Haapaniemi et al., 2018). It is therefore possible that successfully edited cells may not survive to a clonal state, reducing the overall editing efficiency of the CRISPR experiment. To assess whether the enriched environment afforded by our single-cell cloning method could reduce the rate of cell death, we measured the frequency of CRISPR-mediated editing events before and after FACS. We reasoned that equivalence in the number of edits at each of these time points would demonstrate that our method supports cell survival in the face of both (1) DSB, and (2) cell singularisation. The rate of *HPRT* and *FOXP2* editing in the HDFn/HDFa and MICCNi002-A cell lines, respectively, prior to plating was estimated by measuring the NHEJ frequency via a T7EN1 assay in a pool of FACS-sorted positively transfected cells (>5000 cells). We observed that 48.6 to 57.5% of the transfected cells possessed catalytically-induced edits in both the intronic and exonic target regions (**Figure 3A**). To confirm the persistence of the edited cells over the cloning period, we conducted Sanger sequencing of 50 single-cell clones expanded in a 96-well format. Comparable rates of editing in the order 50.0 to 65.0% were observed (**Figure 3B**), indicative of our protocol’s capacity to maintain successfully CRISPR-edited single cells into established clones. We have also measured the editing efficiency of *FOXP2* in the MICCNi002-A cell line using 4D Lonza Nucleofector system to assess whether direct delivery of CRISPR complex to cell nucleus can impact editing frequency. We observed an editing efficiency of 18.66% using nucleofection compared to 56.2% for respective lipofection method (**Supplementary Figure 1**).

**Figure 3.**
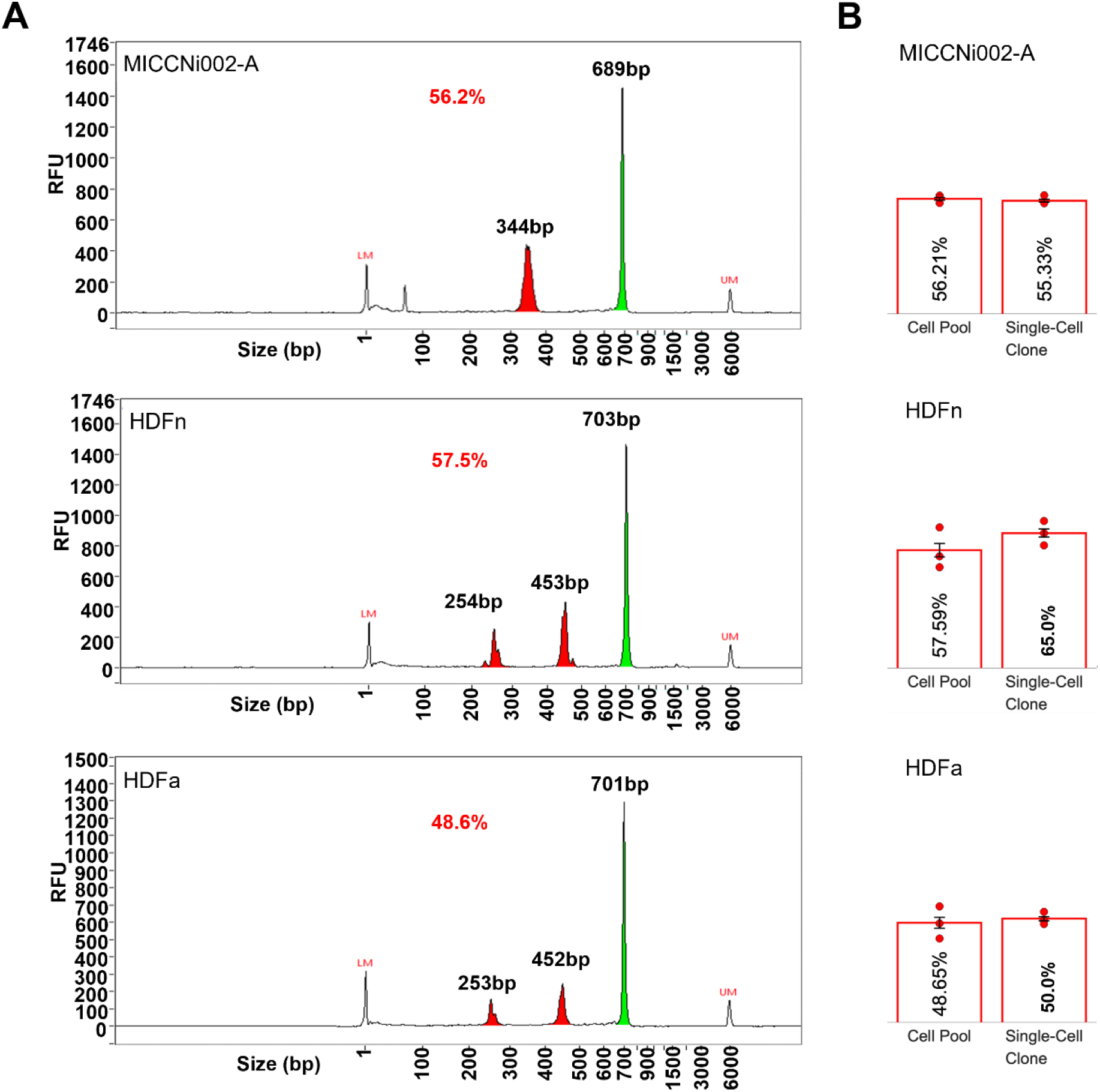
Our method demonstrates successful genome editing. **(A)** Electropherograms of T7EN1 digestion on the pool of FACS-sorted positively transfected cells reveals an editing frequency in the range of 48.6 to 57.5% in the targeted loci. The green (full length) and red (cleaved fragments) peaks are colour-coded for easy identification. RFU = Relative Fluorescence Unit; LM = lower marker; UM = upper marker. **(B)** Editing frequency measured from 50 post transfected single-cell clones (using Sanger sequencing) shows a range of 50.0 to 65.0% which is consistent to their respective cell pool measured by T7EN1 assay. Results are shown as mean of three independent experiments ± SEM.

### The application of our method in HDR editing has successfully resulted in desired knocked-in homogenous iPSC clone

To examine the applicability of our method in HDR, we have performed a knock-in experiment to edit a functionally predicted SNP (rs704074; A/G) in strong linkage disequilibrium with attention deficit hyperactivity disorder Genome wide associated risk variant (Demontis et al., 2019). The MICCNi002-A cell (AA genotype) was transfected with CRISPR-Cas9 complex together with a donor molecule containing the G allele at the SNP site and a T in the PAM site to prevent recutting of the Cas9 enzyme. high-resolution melting analysis (HRMA) screening of 50 positively transfected single-cell clones for the SNP knock-in edit identified 5 with successfully knocked-in variant. Sanger sequencing confirmed the heterozygosity (A/G) genotype, demonstrating a success rate of 10.0% (Pham et al., 2020) (**Figure 4**).

**Figure 4.**
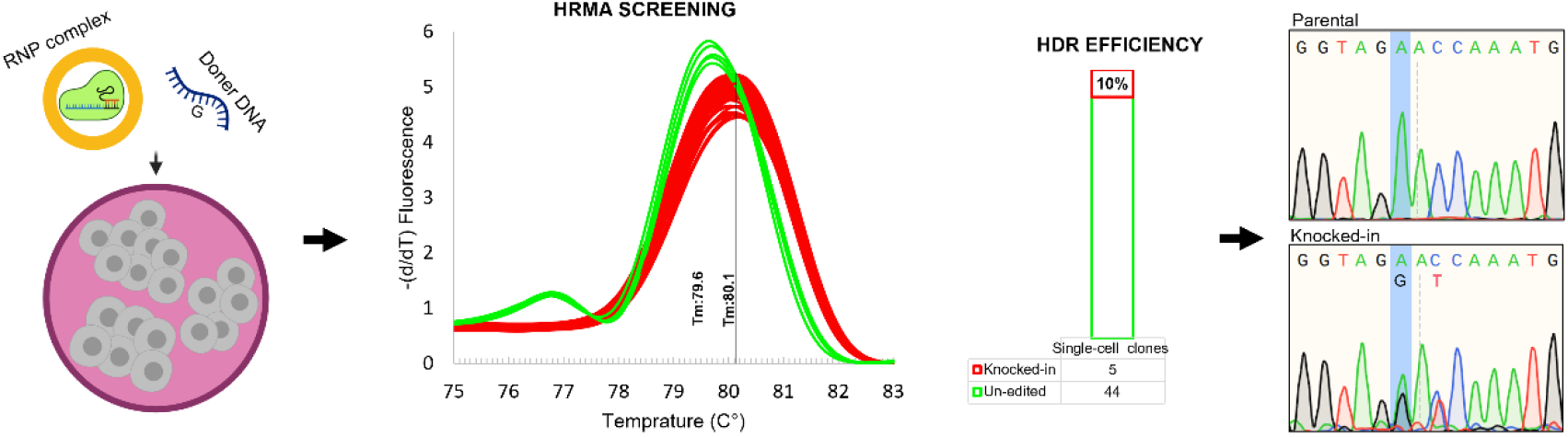
Application of our method in HDR shows the desired editing achieved in two weeks. The MICCNi002-A cell (AA genotype) were transfected with the RNP CRISPR-Cas9 complex targeting rs704074 (A/G) mapped to DUSP*6* gene using the donor molecule containing the G allele. The positively transfected single-cell clones were subjected to HRMA screening to identify the knocked-in clones. Melt curves are normalised using the ratio between fluorescence and temperature variability (-d/dT). Of 50 screened clones, five were identified with different melting temperature (Tm) representing a successful HDR of 10%. This was also confirmed by DNA Sanger sequencing.

### Edited cells show sustained genomic stability, pluripotency, and differentiation potential, as well as lower off-target editing effects

To determine whether the CRISPR-Cas9 genome editing led to complex chromosomal aberrations, we conducted karyotyping to reveal the correct shape, size, and number of chromosomes in the edited cell lines relative to the parental lines (**Supplementary Figure 2**).

To ensure that the edited cell lines retain their utility for disease modelling, we examined the pluripotency and differentiation potential of the edited single-cell clones. Immunostaining for pluripotency markers OCT4, TRA-1-60, SOX2 and SSEA4 in the edited cell lines showed expected expression (**Supplementary Figure 3**), while teratoma testing demonstrated retention of the differentiation potential of the lines into all three germ layers (**Supplementary Figure 4**).

Although the CRISPR-Cas9 complex can generate undesired off-target edits throughout the genome, this risk is reduced with the use of non-integrative CRISPR delivery systems, such as the RNPs applied here. The short half-life of RNPs within the cells further mitigates off-target cutting (Kim et al., 2014). To measure this in our method, we first performed comparative genomic hybridization analysis with a resolution of 200kb between the edited and parental cell lines, to confirm the absence of any major non-target genomic changes. SNPduo analysis (Roberson and Pevsner., 2009) demonstrated that the edited and parental cell lines are identical (**Supplementary Figure S5**). Moreover, there is no evidence of 20q11.21 copy number gains which can protects cells against apoptosis. Further, we Sanger sequenced the top five off-target regions in the pool of FACS-sorted positively transfected cells (>5000 cells) along with their respective un-edited parental lines. Analysis of the sequencing data via the Inference of CRISPR Edits (ICE) web tool (Synthego) and Decomposition tracking of indels revealed no off-target editing to the prenatal lines (**Supplementary Table S3**) (Conant et al., 2022).

Collectively, these findings suggest that the RNP-lipid delivery of the CRISPR complex, together with the provision of Laminin-521, StemFlex medium, and CloneR supplement to individual transfected iPSCs, can significantly improve the establishment and expansion of genetically homogeneous single-cell clones, while maintaining a high editing efficacy. Figure 5 shows our workflow for CRISPR engineering of iPSCs and expansion of single cell clones in 96-well plate format.

**Figure 5.**
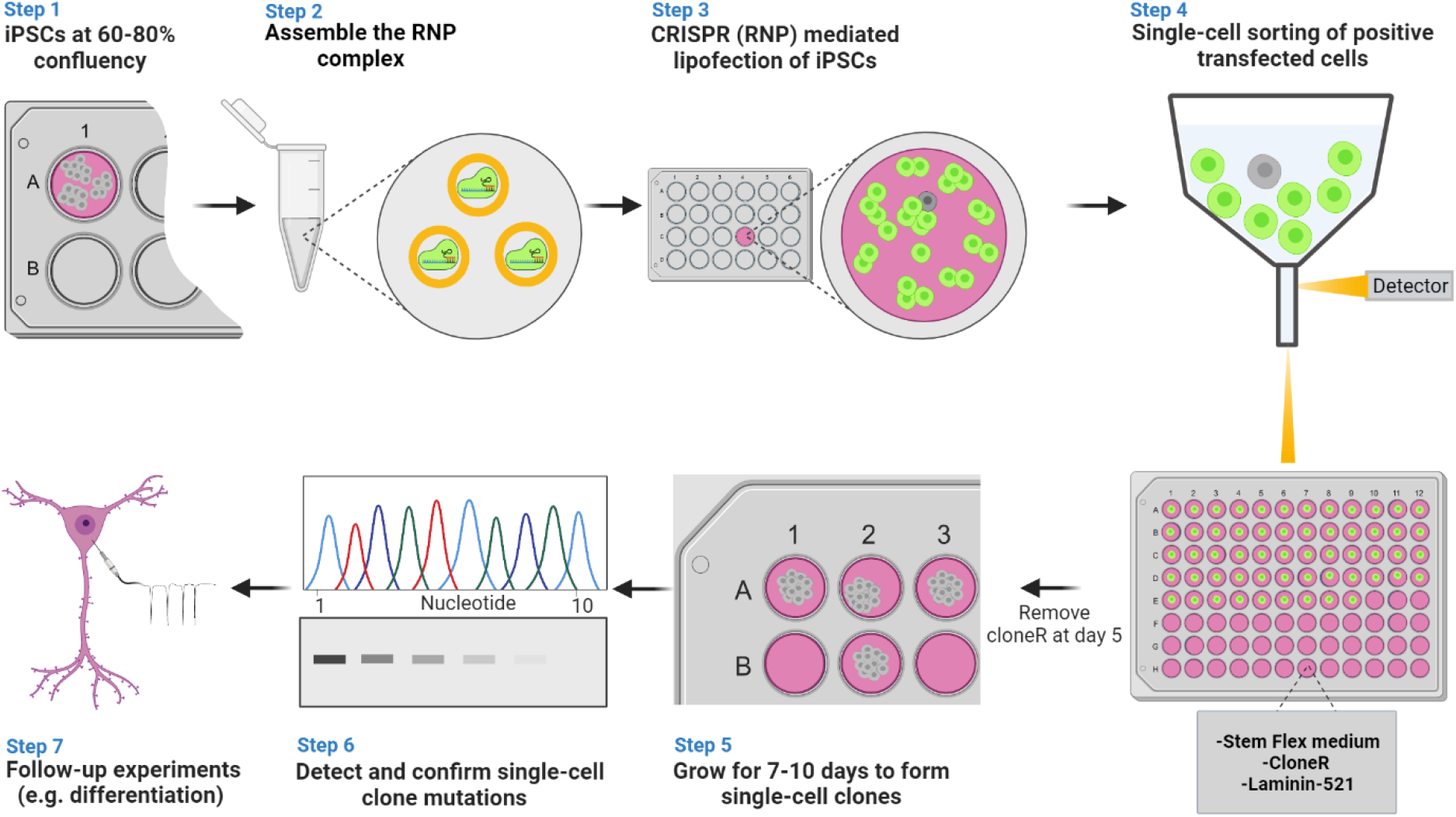
Schematic workflow for our genetically homogenous CRISPRed-iPSC single cell cloning. **Step 1:** iPSCs are grown to 60-80% confluency. **Step 2:** The ribonucleoprotein complex (RNP) consisting of gRNA:tracrRNA duplex and Cas9 protein is formed and transfected within the small clusters of iPSC cells using Lipofectamine Stem Transfection Reagent **(Step 3)**. **Step 4:** Following 48 hours of incubation, positively transfected cells are single-cell sorted in a rich environment containing StemFlex medium and CloneR supplement in wells of 96-well plate coated with Laminin-521. **Step 5:** Single-cells are grown for 7-10 days until they form a homogeneous single-cell clone and reach sufficient confluency for expansion. **Step 6:** A portion of the clones are selected for detection and confirmation of the desired mutations (e.g., Sanger sequencing, western blotting) and the remainder utilised for follow-up phenotyping experiments **(Step 7)**. Figure created with BioRender.com.

## DISCUSSION

Here we describe a simple and efficient method for genetically homogeneous single-cell cloning of iPSCs following transfection with CRISPR-editing machinery. The most significant benefit of our method is the substantially higher survival rate of up to 70.1% of transfected iPSC clones. This high survival rate is largely due to our efforts to diminish the toxicity of the CRISPR transfection process on cell survival, as well as provision of an enriched environment for the establishment and expansion of isolated CRISPRed iPSCs. The former was fulfilled via the gentle lipidic delivery of RNP complexes. The alternative delivery method of electroporation, although common, is an incredibly invasive method, requiring both electrical shock to the cells and cell singularisation. These result in low post-treatment recovery of electroporated cells and increased risk of apoptosis (Stewart et al., 2018). When applied with DNA-based vectors (i.e., plasmid, viral particles), the risk of cellular toxicity is compounded (Shimokawa et al., 2000; Stacey et al., 1993; Van De Parre et al., 2005). For instance, combined electroporation-plasmid protocols have reported up to 95% cell death following transfection (Li et al., 2018), limiting the pool of potentially edited cells available for further analysis. By contrast, application of less toxic protein-based delivery systems (RNPs) together with gentle lipofectamine reagents, require that cells only be dissociated into small clumps of 3-10 cells, significantly improving cell survival post-transfection (Chen et al., 2010).

The supportive conditions for homogeneous single-cell clonal formation were achieved through gradual optimisation of the three major elements fortifying individual CRISPRed iPSC’s viability. Through this, we demonstrated that the combination of StemFlex medium, CloneR supplementation, and Laminin-521 surface martix greatly enhances the survival of isolated transfected iPSCs, permitting them to expand into genetically homogeneous clones. Laminin-521 as a key cell adhesion molecule is predominantly found in the stem cell niches, indicating the importance of this protein for the growth of stem cells, and has been reported to improve clonogenicity of human hPSCs by up to 20% (Rodin et al., 2014). When used in tandem with Stemflex and cloneR, these reagents aid single-cell survival by establishing a supportive protein scaffolding prior to plating, while simultaneously alleviating the stressful state of iPSC singularisation once plated, thereby enhancing clonal expansion (Daniszewski et al., 2018). Although application of other small molecule cocktails has been reported to increase the clonal rate up to 80%, the impact of CRISPR machinery toxicity on single-cell clone survival has not been considered (Chen et al., 2021; Chen and Pruett-Miller, 2018; Tristan et al., 2022). This is being reflected in our experiment by the approximately 20% increase of the non-CRISPRed clonal survival rate, leading up to 90% clonogenicity. Such high viability in our method was achieved without the use of feeder layer cells, mitigating the risk of retroviral contamination, and ensuring that any observable difference in cell morphology/function between lines is the product of isogeneic variation (Cobo et al., 2008). Importantly, the high single-cell cloning described is incredibly efficient—achieving genetically homogeneous clones in between 7 to 10 days. This suggests that our protocol is suitable for rapid upscaling of gene function analysis.

Simultaneous application of our cloning method with the use of CRISPR RNPs affords additional benefits to the CRISPR editing process. Their non-integrative mode of action and rapid degradation in the intracellular environment reduces recutting of the target locus, thereby improving editing efficiency (Okamoto et al., 2019; Vakulskas et al., 2018). As demonstrated, application of our cloning method together with the lipidic delivery of CRISPR RNPs resulted in an NHEJ editing frequency of between 50.0 to 65.0% of the surviving single iPSC clones. This result coincided a significantly high transfection efficiency of >97%. Moreover, the consistency of the editing frequency prior to and post single-cell cloning validated that the toxicity of the CRISPR complex is minimised in our cloning method. It is of note that editing efficiency and cell survival is locus and cell line dependant (Cahan and Daley, 2013). Less successful viability and editing may be observed when targeting essential or loss-of-function intolerant genes, and may be influenced by variability in the genetic background between cell lines. We expect, however, that the chance of cell survival with our method is likely to be relatively enhanced in these instances. This can be inferred by the persistence of both our intronically and exonically edited iPSC clones, suggesting that the combination of reagents described supports cell growth irrespective of target gene.

Integrating our genetically homogenous single-cell cloning approach with the single nucleotide polymorphism HDR system can help to avoid standard practices of cellular expansion in mixed cell populations (Deneault et al., 2019; Miyaoka *et al*., 2014; Schrode et al., 2019). This not only negates the need for labour-intensive and time-consuming enrichment steps, but also enhances the odds of capturing clones with the intended on-target edit. It also has the capacity to be combined with epigenome editing approaches such as CRISPRi and CRISPRa, ensuring homogeneous transcription of the target gene(s) within a cell population.

In brief, our method addresses a key challenge posed by the application of CRISPR-Cas9 technology in human derived iPSCs: achieving high rates of CRISPRed single-cell clone survival while maintaining monoclonality. Our quick and simple cloning method supports the expansion of transfected single-cell clones, thereby improving homogeneity of the resultant cell lines for valid isogenic cell line phenotypic comparison. We anticipate that our approach will facilitate stem cell modelling of complex genetic conditions, bridging the gap between identification of genomic risk loci and their associated functional biology.

## EXPERIMENTAL PROCEDURES

### Corresponding Author

Further information and requests concerning our data should be addressed to corresponding authors, Ziarih Hawi and/or Atefeh Namipashaki

### Materials Availability

This study did not generate new unique materials/reagents.

### Data availability

All data generated and analysed during this study are included in the article and supporting files.

### Guide RNA (gRNA) and primer design

The Alt-R CRISPR-Cas9 CRISPR RNA (crRNA) selection tool from Integrated DNA Technology (IDT) (https://sg.idtdna.com/site/order/designtool/index/CRISPR_PREDESIGN) was used to design donor molecule and gRNAs targeting exon 2 of the Forkhead Box P2 (*FOXP2*) gene and DUSP6 Gene 3 prime UTR variant. Predesigned Alt-R CRISPR-Cas9 HPRT Positive Control crRNA targeting intron 6-7 of the human Hypoxanthine Phosphoribosyltransferase 1 (*HPRT*) gene and the non-targeting Alt-R CRISPR-Cas9 Negative Control crRNA (IDT, Cat#1079132) was purchased from IDT. Detailed sequences are provided in Supplementary Table S1. Primer sequences used to amplify the targeted and the top five potential off-target regions were designed using primer3plus (https://www.bioinformatics.nl/cgi-bin/primer3plus/primer3plus.cgi) (Supplementary Table S2-3).

### Pre-transfection iPSCs culture

Three iPSC cell lines were used in this study. These consisted of the peripheral blood mononuclear cells (PBMC)-derived MICCNi002-A (Tong et al., 2019), and Fibroblast derived HDFn and HDFa. All lines were generated in-house using the CytoTune™-iPS 2.0 Sendai Reprogramming Kit (Invitrogen, Cat#A16517). Human Dermal Fibroblasts, neonatal (HDFn) (Cat#C0045C) and Human Dermal Fibroblasts, adult (HDFa) (Cat#C0135C) were purchased commercially from Thermo Fisher Scientific. All iPSC lines were seeded on Vitronectin (Thermo Fisher Scientific, Cat#A14700)-coated plates and were maintained in Essential 8 Medium (Gibco, Cat#A1517001) supplemented with RevitaCell (Gibco, Cat# A2644501) and 1X Penicillin-Streptomycin (Gibco, Cat#15140122) for the first 24 hours. Media were changed every day with Essential 8 medium without RevitaCell until 60-80% confluency was reached, allowing for up to three passages for cells to adapt and reach a healthy state with a minimum differentiation rate.

### Construction of the RNP CRISPR transfection complex

One microliter (µL) of ALT-R CRISPR-CAS9 gRNA (100 µM) and Alt-R CRISPR-Cas9 tracrRNA (100 µM) conjugated with ATTO™ 550 (Cat#1075928) were added to 98 µL of Nuclease-free Duplex Buffer (Cat#11-01-03-01) to form the gRNA:tracrRNA duplex. The ATTO-550 is a fluorescent compound with an excitation and emission peaks at 553 nm and 575 nm respectively. The mixture was heated at 95°C for 5 minutes and allowed to cool for 30 minutes at room temperature. To assemble the RNP complex, 6 µL of an equimolar (1 µM) amount of diluted Cas9 Nuclease (Cat#1081058) and gRNA:tracrRNA duplex were mixed with 13 µL of Opti-MEM Medium (Thermo Fisher Scientific, Cat#51985091) and incubated for 5 minutes at room temperature. All reagents were purchased from IDT unless otherwise specified.

### Lipid delivery of the CRISPR complex

Two microlitres of Lipofectamine Stem Transfection Reagent (Invitrogen, Cat#STEM00015) and 23 µL Opti-MEM medium were mixed and added to the 25 µL of RNP complex. The total CRISPR-Cas9 transfection complex (50 µL) was incubated for 20 minutes at room temperature prior to addition to the iPSCs. At 60-80% confluency, Versene solution (Gibco, Cat# 15040066) was added to the iPSC cell culture and was subsequently aspirated following 5 minutes of incubation. Essential 8 medium supplemented with RevitaCell without antibiotic was added immediately to the cells, pipetting up and down several times to dissociate them into clusters of 3-10 cells. The cells were then counted and adjusted to 300,000 cell/ml. One hundred and fifty thousand cells were added to 50 µL of transfection complex pre-plated in a single well of a 24-well plate coated with Vitronectin containing Essential 8 medium with RevitaCell supplement. For the knock-in experiment, 1.0 µM of the donor molecule containing the G allele and 0.3uL of HDR enhancer (IDT, Cat#10007910) were added to the transfection complex.

### Electroporation of the CRISPR complex

For electroporation, 3.2 µL of the RNP complex was mixed with 16.4 µL of the Lonza P3 electroporation solution. The total solution was added to a cell pellet containing 250,000 singularised iPSCs (using TrypLE™ Select Enzyme [Gibco, Cat# A1217701]). The cell suspension was transferred to a well of 20 µL Nucleocuvette strip and subjected to electroporation using the CB-150 program in a 4D Lonza Nucleofector system. Following 10 minutes of recovery, the transfected cells were added to a single well of a 24-well plate coated with Vitronectin containing Essential 8 medium with RevitaCell supplement.

### Sorting the positively transfected cells

Transfected cells were dissociated into single cells with Accutase (STEMCELL Technologies, Cat#07920) and centrifuged for 4 minutes at 200 x g. The cell palette was resuspended in 300 µL of FACS medium prepared with 4 mL of DMEM/F-12, HEPES, no phenol red (Gibco, Cat# 11039021), 0.5 mL of CloneR supplement (STEMCELL Technologies, Cat# 05889), 0.5 mL of E8 supplement (Gibco, Cat#A15171-01), 50 µL of (100X) Penicillin-Streptomycin (Gibco, Cat#15140122), 5 µL of DAPI (Cell Signaling Technology, Cat#4083S), and 2 µL of 0.5 M EDTA (Invitrogen, Cat# 15575020) and passed through a Falcon 5 mL tube with a cell strainer cap (Corning Cat# 352235). This supplemented FACS medium enhances the survival of singularised iPSCs for the duration of sorting. Utilising a BD Influx Cell Sorter (BD Biosciences), the top 40% of ATTO-550 positive cells were sorted as single-cells into individual wells of two 96-well plates with different combination of CellAdhere Laminin-521 (STEMCELL Technologies, Cat#77004, (2 μg/cm^2^)) or Vitronectin (0.5 μg/cm^2^), StemFlex medium (Gibco, Cat#A3349401) or Essential 8 Flex Medium (Gibco, Cat# A2858501) and supplementation of RevitaCell or CloneR and incubated at 37°C for 48 hours. The remaining positive cells were bulk sorted in a single well of a Vitronectin coated 6-well plate containing Essential 8 medium and ROCK inhibitor (STEMCELL Technologies, Cat#72304) for the purpose of measuring editing efficiency.

### Post-FACS single-cell iPSC cell culture

Following 48 hours of incubation, a full CloneR-supplemented media change was performed. At day three, 25% of the initial seed volume of cloning media were added to each well. At day four, a full media without CloneR supplement was performed and repeated every other day until the single-cell clones had emerged (∼60-80% confluency) for further analysis. On day 10, the total number of single-cell clones from each cell line were counted to obtain single-cell clone survival rate.

### Measuring editing frequency

Total genomic DNA from the pool of FACS-sorted positively transfected cells (>5000 cells) for HDFn and HDFa and MICCNi002-A were extracted using PureLink Genomic DNA Mini Kit (Invitrogen, Cat#K182001). The *HPRT* and *FOXP2* target regions were amplified using PCRBIO HiFi Polymerase (PCR Biosystems, Cat#PB10.41-02) with primers listed in Supplementary Table S2.

The PCR products were digested with T7 endonuclease 1 enzyme using the Alt-R Genome Editing Detection Kit (Integrated DNA Technologies, Cat#1075931). The cleaved DNA fragments were run on an Agilent Fragment Analyzer system and analysed with the ProSize Software (Agilent Technologies) to measure the size of and quantify the fragments. The frequency of NHEJ was calculated using the below formula, as described by Agilent Technologies:

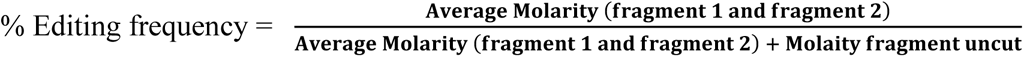

To confirm the achieved editing efficiency following single-cell cloning, 50 single-cell clones from each cell line were expanded in 6-well plate. DNA was extracted and amplified as described above. The targeted PCR amplicons of *HPRT* and *FOXP2* from each clone were subsequently Sanger sequenced. The percentage of edited clones were calculated as the number of clones detected to possess at least one bp change relative to the total number of sequenced clones.

### HRMA screening

Isolated genomic DNA (10ng) from 50 single-cell clone were amplified LightCycler 480 SYBR Green I Master and were subjected to melt curve analysis to screen for the edited clones.

### Karyotyping

To discern whether the CRISPR-editing protocol induced chromosomal aberrations such as deletions, duplications, translocations, inversions, the edited single-cell clones from each cell line were externally Karyotyped and analysed using standard methods at Monash Health Pathology, Australia.

### Immunofluorescence

Pluripotency of the CRISPR-Cas9 edited single-cell clones was evaluated via immunostaining of OCT4, TRA-1-60, SOX2 and SSEA4 markers using the Pluripotent Stem Cell 4-Marker Immunocytochemistry Kit (Invitrogen, Cat#3A24881). The culturing media were aspirated, and the cells were incubated in fixative solution for 15 minutes, followed by 30 minutes of blocking, then an additional period of incubation with the primary antibodies for 3 hours at 4°C. The cells were subsequently washed and further incubated with the appropriate secondary antibodies for 1 hour at room temperature. This was followed by further washing steps, with the last wash containing NucBlue Fixed Cell Stain (DAPI). The cells were immediately imaged with the EVOS M5000 Imaging System (Thermo Fisher Scientific, Cat# AMF5000).

### Teratoma formation assay

Approximately 1 x 10^6^ iPSCs from each of the three edited cell lines were injected into severe combined immunodeficiency disease (SCID) mice. Six weeks later, 1-2 cm diameter teratomas formed and were subsequently hematoxylin and eosin stained at the Monash Histology Platform to examine their ability to differentiate into each of endoderm, ectoderm, and the mesoderm germ layers.

### Comparative genomic hybridization array

To scan the entire genome for genetic imbalance (e.g., copy number gains and losses), micro-array based comparative genomic hybridization of the CRISPR edited lines relative to wildtype genome was conducted at the Victorian Clinical Genetics Services (Melbourne, Australia).

### Analysis of off-target mutation using ICE

The top five off-target genomic regions of the transfected cell pool and the parental cell lines were amplified, and Sanger sequenced. Results were compared to the wild type using ICE: a Synthego web tool (https://www.synthego.com/products/bioinformatics/crispr-analysis) which uses Sanger sequencing data to analyse the levels of indels caused by CRISPR editing (Supplementary Table S3).

### Statistical information

The samples for measurement of cell survival and editing frequency were all biological replicates (independent transfections); calculated by the mean of independent experiments ± standard error of mean (SEM). Statistical differences for cell viability between the lipofection and electroporation method were evaluated using Student’s *t* tests.

## Acknowledgements

This work has been supported by Project Grant funding from the National Health and Medical Research Council (NHMRC) of Australia to ZH and MAB (APP1146644).

## Author contributions

A.N, X.L and Z.H designed the idea and planned the experiments. A.N performed all experiments except of teratoma formation assay which was performed by SM.L, G.S and *FOXP2* CRISPR experiment which was performed by K.A and A.N. Z.H, M.A.B, J.M.P and M.J.H developed the theory, contributed to the interpretation of the results, and oversaw the overall direction. The manuscript was prepared by A.N, Z.H and K.P. All authors contributed to comments and editing.

## Ethics declaration

The teratoma assays for human stem cells and reprogrammed cells have been performed under the Monash University Animal Ethics Research number ERM # 26424.

## References

1. Byrne, S.M., Mali, P., and Church, G.M. (2014). Genome editing in human stem cells. In Methods in enzymology, (Elsevier), pp. 119–138. 10.1016/B978-0-12-801185-0.00006-4

2. Cahan, P., and Daley, G.Q. (2013). Origins and implications of pluripotent stem cell variability and heterogeneity. Nat Rev Mol Cell Biol 14, 357–368. 10.1038/nrm3584

3. Chang, C.Y., Ting, H.C., Su, H.L., and Jeng, J.R. (2018). Combining Induced Pluripotent Stem Cells and Genome Editing Technologies for Clinical Applications. Cell Transplant 27, 379–392. 10.1177/0963689718754560

4. Chen, G., Hou, Z., Gulbranson, D.R., and Thomson, J.A. (2010). Actin-myosin contractility is responsible for the reduced viability of dissociated human embryonic stem cells. Cell Stem Cell 7, 240–248. 10.1016/j.stem.2010.06.017

5. Chen, Y., Tristan, C.A., Chen, L., Jovanovic, V.M., Malley, C., Chu, P.H., Ryu, S., Deng, T., Ormanoglu, P., Tao, D., et al. (2021). A versatile polypharmacology platform promotes cytoprotection and viability of human pluripotent and differentiated cells. Nat Methods 18, 528–541. 10.1038/s41592-021-01126-2

6. Chen, Y.H., and Pruett-Miller, S.M. (2018). Improving single-cell cloning workflow for gene editing in human pluripotent stem cells. Stem Cell Res 31, 186–192. 10.1016/j.scr.2018.08.003

7. Cobo, F., Navarro, J.M., Herrera, M.I., Vivo, A., Porcel, D., Hernandez, C., Jurado, M., Garcia-Castro, J., and Menendez, P. (2008). Electron microscopy reveals the presence of viruses in mouse embryonic fibroblasts but neither in human embryonic fibroblasts nor in human mesenchymal cells used for hESC maintenance: toward an implementation of microbiological quality assurance program in stem cell banks. Cloning Stem Cells 10, 65–74. 10.1089/clo.2007.0020

8. Conant, D., Hsiau, T., Rossi, N., Oki, J., Maures, T., Waite, K., Yang, J., Joshi, S., Kelso, R., Holden, K., et al. (2022). Inference of CRISPR Edits from Sanger Trace Data. CRISPR J 5, 123–130. 10.1089/crispr.2021.0113

9. Daniszewski, M., Nguyen, Q., Chy, H.S., Singh, V., Crombie, D.E., Kulkarni, T., Liang, H.H., Sivakumaran, P., Lidgerwood, G.E., Hernandez, D., et al. (2018). Single-Cell Profiling Identifies Key Pathways Expressed by iPSCs Cultured in Different Commercial Media. iScience 7, 30–39. 10.1016/j.isci.2018.08.016

10. Das, D., Feuer, K., Wahbeh, M., and Avramopoulos, D. (2020). Modeling Psychiatric Disorder Biology with Stem Cells. Curr Psychiatry Rep 22, 24. 10.1007/s11920-020-01148-1

11. Demontis, D., Walters, G.B., Athanasiadis, G., Walters, R., Therrien, K., Nielsen, T.T., Farajzadeh, L., Voloudakis, G., Bendl, J., Zeng, B. and Zhang, W., 2023. Genome-wide analyses of ADHD identify 27 risk loci, refine the genetic architecture and implicate several cognitive domains. Nature genetics, 55(2), pp.198–208. 10.1038/s41588-022-01285-8

12. Deneault, E., Faheem, M., White, S.H., Rodrigues, D.C., Sun, S., Wei, W., Piekna, A., Thompson, T., Howe, J.L., Chalil, L., et al. (2019). CNTN5(-)(/+)or EHMT2(-)(/+)human iPSC-derived neurons from individuals with autism develop hyperactive neuronal networks. Elife 8. 10.7554/elife.40092

13. Gallagher, M.D., and Chen-Plotkin, A.S. (2018). The Post-GWAS Era: From Association to Function. Am J Hum Genet 102, 717–730. 10.1016/j.ajhg.2018.04.002

14. Geng, B.C., Choi, K.H., Wang, S.Z., Chen, P., Pan, X.D., Dong, N.G., Ko, J.K., and Zhu, H. (2020). A simple, quick, and efficient CRISPR/Cas9 genome editing method for human induced pluripotent stem cells. Acta Pharmacol Sin 41, 1427–1432. 10.1038/s41401-020-0452-0

15. Haapaniemi, E., Botla, S., Persson, J., Schmierer, B., and Taipale, J. (2018). CRISPR-Cas9 genome editing induces a p53-mediated DNA damage response. Nat Med 24, 927–930. 10.1038/s41591-018-0049-z

16. Hamm, A., Krott, N., Breibach, I., Blindt, R. and Bosserhoff, A.K. (2002). Efficient transfection method for primary cells. Tissue Eng, 8, 235–245. 10.1089/107632702753725003.

17. Hoffmann, A., Ziller, M., and Spengler, D. (2020). Focus on Causality in ESC/iPSC-Based Modeling of Psychiatric Disorders. Cells 9. 10.3390/cells9020366

18. Ihry, R.J., Worringer, K.A., Salick, M.R., Frias, E., Ho, D., Theriault, K., Kommineni, S., Chen, J., Sondey, M., Ye, C., et al. (2018). p53 inhibits CRISPR-Cas9 engineering in human pluripotent stem cells. Nat Med 24, 939–946. 10.1038/s41591-018-0050-6

19. Ikeda, K., Uchida, N., Nishimura, T., White, J., Martin, R.M., Nakauchi, H., Sebastiano, V., Weinberg, K.I., and Porteus, M.H. (2018). Efficient scarless genome editing in human pluripotent stem cells. Nat Methods 15, 1045–1047. 10.1038/s41592-018-0212-y

20. Kampmann, M. (2020). CRISPR-based functional genomics for neurological disease. Nat Rev Neurol 16, 465–480. 10.1038/s41582-020-0373-z

21. Kim, S., Kim, D., Cho, S.W., Kim, J., and Kim, J.S. (2014). Highly efficient RNA-guided genome editing in human cells via delivery of purified Cas9 ribonucleoproteins. Genome Res 24, 1012–1019. 10.1101/gr.171322.113

22. Kim, T.K., and Eberwine, J.H. (2010). Mammalian cell transfection: the present and the future. Anal Bioanal Chem 397, 3173–3178. 10.1007/s00216-010-3821-6

23. Kwart, D., Paquet, D., Teo, S., and Tessier-Lavigne, M. (2017). Precise and efficient scarless genome editing in stem cells using CORRECT. Nat Protoc 12, 329–354. 10.1038/nprot.2016.171

24. Li, H.L., Gee, P., Ishida, K., and Hotta, A. (2016). Efficient genomic correction methods in human iPS cells using CRISPR-Cas9 system. Methods 101, 27–35. 10.1016/j.ymeth.2015.10.015

25. Li, X., Krawetz, R., Liu, S., Meng, G., and Rancourt, D.E. (2009). ROCK inhibitor improves survival of cryopreserved serum/feeder-free single human embryonic stem cells. Hum Reprod 10.1093/humrep/den404

26. Li, X.L., Li, G.H., Fu, J., Fu, Y.W., Zhang, L., Chen, W., Arakaki, C., Zhang, J.P., Wen, W., Zhao, M., et al. (2018). Highly efficient genome editing via CRISPR-Cas9 in human pluripotent stem cells is achieved by transient BCL-XL overexpression. Nucleic Acids Res 46, 10.1093/nar/gky804

27. Lino, C.A., Harper, J.C., Carney, J.P., and Timlin, J.A. (2018). Delivering CRISPR: a review of the challenges and approaches. Drug Deliv 25, 1234–1257. 10.1080/10717544.2018.1474964

28. Matos, M.R., Ho, S.M., Schrode, N., and Brennand, K.J. (2020). Integration of CRISPR-engineering and hiPSC-based models of psychiatric genomics. Mol Cell Neurosci 107, 103532. 10.1016/j.mcn.2020.103532

29. Merkle, F.T., Neuhausser, W.M., Santos, D., Valen, E., Gagnon, J.A., Maas, K., Sandoe, J., Schier, A.F., and Eggan, K. (2015). Efficient CRISPR-Cas9-mediated generation of knockin human pluripotent stem cells lacking undesired mutations at the targeted locus. Cell Rep 11, 875–883. 10.1016/j.celrep.2015.04.007

30. Mianne, J., Bourguignon, C., Nguyen Van, C., Fieldes, M., Nasri, A., Assou, S., and De Vos, J. (2020). Pipeline for the Generation and Characterization of Transgenic Human Pluripotent Stem Cells Using the CRISPR/Cas9 Technology. Cells 9. 10.3390/cells9051312

31. Miyaoka, Y., Chan, A.H., Judge, L.M., Yoo, J., Huang, M., Nguyen, T.D., Lizarraga, P.P., So, P.L., and Conklin, B.R. (2014). Isolation of single-base genome-edited human iPS cells without antibiotic selection. Nat Methods 11, 291–293. 10.1038/nmeth.2840

32. Ohgushi, M., Matsumura, M., Eiraku, M., Murakami, K., Aramaki, T., Nishiyama, A., Muguruma, K., Nakano, T., Suga, H., Ueno, M., et al. (2010). Molecular pathway and cell state responsible for dissociation-induced apoptosis in human pluripotent stem cells. Cell Stem Cell 7, 225–239. 10.1016/j.stem.2010.06.018

33. Okamoto, S., Amaishi, Y., Maki, I., Enoki, T., and Mineno, J. (2019). Highly efficient genome editing for single-base substitutions using optimized ssODNs with Cas9-RNPs. Sci Rep 9, 4811. 10.1038/s41598-019-41121-4

34. Pham, Q.T., Raad, S., Mangahas, C.L., M’callum, M.A., Raggi, C. and Paganelli, M. (2020). High-throughput assessment of mutations generated by genome editing in induced pluripotent stem cells by high-resolution melting analysis. Cytotherapy, 22, 536–542. 10.1016/j.jcyt.2020.06.008

35. Roberson, E.D. and Pevsner, J. (2009). Visualization of shared genomic regions and meiotic recombination in high-density SNP data. PloS one 4, e6711. 10.1371/journal.pone.0006711

36. Rodin, S., Antonsson, L., Hovatta, O., and Tryggvason, K. (2014). Monolayer culturing and cloning of human pluripotent stem cells on laminin-521-based matrices under xeno-free and chemically defined conditions. Nat Protoc 9, 2354–2368. 10.1038/nprot.2014.159

37. Sander, J.D., and Joung, J.K. (2014). CRISPR-Cas systems for editing, regulating and targeting genomes. Nat Biotechnol 32, 347–355. 10.1038/nbt.2842

38. Schrode, N., Ho, S.M., Yamamuro, K., Dobbyn, A., Huckins, L., Matos, M.R., Cheng, E., Deans, P.J.M., Flaherty, E., Barretto, N., et al. (2019). Synergistic effects of common schizophrenia risk variants. Nat Genet 51, 1475–1485. 10.1038/s41588-019-0497-5

39. Sharifi Tabar, M., Hesaraki, M., Esfandiari, F., Sahraneshin Samani, F., Vakilian, H., and Baharvand, H. (2015). Evaluating Electroporation and Lipofectamine Approaches for Transient and Stable Transgene Expressions in Human Fibroblasts and Embryonic Stem Cells. Cell J 17, 438–450. 10.22074/cellj.2015.5

40. Shimokawa, T., Okumura, K., and Ra, C. (2000). DNA induces apoptosis in electroporated human promonocytic cell line U937. Biochem Biophys Res Commun 270, 94–99. 10.1006/bbrc.2000.2388

41. Simkin, D., Papakis, V., Bustos, B.I., Ambrosi, C.M., Ryan, S.J., Baru, V., Williams, L.A., Dempsey, G.T., McManus, O.B., Landers, J.E., et al. (2022). Homozygous might be hemizygous: CRISPR/Cas9 editing in iPSCs results in detrimental on-target defects that escape standard quality controls. Stem Cell Reports 17, 993–1008. 10.1016/j.stemcr.2022.02.008

42. Singh, A.M. (2019). An Efficient Protocol for Single-Cell Cloning Human Pluripotent Stem Cells. Front Cell Dev Biol 7, 11. 10.3389/fcell.2019.00011

43. Soldner, F., Laganiere, J., Cheng, A.W., Hockemeyer, D., Gao, Q., Alagappan, R., Khurana, V., Golbe, L.I., Myers, R.H., Lindquist, S., et al. (2011). Generation of isogenic pluripotent stem cells differing exclusively at two early onset Parkinson point mutations. Cell 146, 318–331. 10.1016/j.cell.2011.06.019

44. Stacey, K.J., Ross, I.L., and Hume, D.A. (1993). Electroporation and DNA-dependent cell death in murine macrophages. Immunol Cell Biol 71 ( *Pt 2*), 75–85. 10.1038/icb.1993.8

45. Stewart, M.P., Langer, R., and Jensen, K.F. (2018). Intracellular Delivery by Membrane Disruption: Mechanisms, Strategies, and Concepts. Chem Rev 118, 7409–7531. 10.1021/acs.chemrev.7b00678

46. Steyer, B., Bu, Q., Cory, E., Jiang, K., Duong, S., Sinha, D., Steltzer, S., Gamm, D., Chang, Q., and Saha, K. (2018). Scarless Genome Editing of Human Pluripotent Stem Cells via Transient Puromycin Selection. Stem Cell Reports 10, 642–654. 10.1016/j.stemcr.2017.12.004

47. Tong, J., Lee, K.M., Liu, X., Nefzger, C.M., Vijayakumar, P., Hawi, Z., Pang, K.C., Parish, C.L., Polo, J.M., and Bellgrove, M.A. (2019). Generation of four iPSC lines from peripheral blood mononuclear cells (PBMCs) of an attention deficit hyperactivity disorder (ADHD) individual and a healthy sibling in an Australia-Caucasian family. Stem Cell Res 34, 101353. 10.1016/j.scr.2018.11.014

48. Tristan, C.A., Hong, H., Jethmalani, Y., Chen, Y., Weber, C., Chu, P.H., Ryu, S., Jovanovic, V.M., Hur, I., Voss, T.C., et al. (2022). Efficient and safe single-cell cloning of human pluripotent stem cells using the CEPT cocktail. Nat Protoc. 10.1038/s41596-022-00753-z

49. Vakulskas, C.A., Dever, D.P., Rettig, G.R., Turk, R., Jacobi, A.M., Collingwood, M.A., Bode, N.M., McNeill, M.S., Yan, S., Camarena, J., et al. (2018). A high-fidelity Cas9 mutant delivered as a ribonucleoprotein complex enables efficient gene editing in human hematopoietic stem and progenitor cells. Nat Med 24, 1216–1224. 10.1038/s41591-018-0137-0

50. Van De Parre, T.J., Martinet, W., Schrijvers, D.M., Herman, A.G., and De Meyer, G.R. (2005). mRNA but not plasmid DNA is efficiently transfected in murine J774A.1 macrophages. Biochem Biophys Res Commun 327, 356–360. 10.1016/j.bbrc.2004.12.027

51. Wang, G., Yang, L., Grishin, D., Rios, X., Ye, L.Y., Hu, Y., Li, K., Zhang, D., Church, G.M., and Pu, W.T. (2017). Efficient, footprint-free human iPSC genome editing by consolidation of Cas9/CRISPR and piggyBac technologies. Nat Protoc 12, 88–103. 10.1038/nprot.2016.152

52. Watanabe, K., Ueno, M., Kamiya, D., Nishiyama, A., Matsumura, M., Wataya, T., Takahashi, J.B., Nishikawa, S., Nishikawa, S., Muguruma, K., and Sasai, Y. (2007). A ROCK inhibitor permits survival of dissociated human embryonic stem cells. Nat Biotechnol 25, 681–686. 10.1038/nbt1310

53. Xu, X., Gao, D., Wang, P., Chen, J., Ruan, J., Xu, J., and Xia, X. (2018). Efficient homology-directed gene editing by CRISPR/Cas9 in human stem and primary cells using tube electroporation. Sci Rep 8, 11649. 10.1038/s41598-018-30227-w

54. Yang, L., Yang, J.L., Byrne, S., Pan, J., and Church, G.M. (2014). CRISPR/Cas9-directed genome editing of cultured cells. Current protocols in molecular biology 107, 31.31.31–31.31.17. 10.1002/0471142727.mb3101s107

55. Yumlu, S., Stumm, J., Bashir, S., Dreyer, A.K., Lisowski, P., Danner, E., and Kuhn, R. (2017). Gene editing and clonal isolation of human induced pluripotent stem cells using CRISPR/Cas9. Methods 121-122, 29-44. 10.1016/j.ymeth.2017.05.009

56. Zizzi, A., Minardi, D., Ciavattini, A., Giantomassi, F., Montironi, R., Muzzonigro, G., Di Primio, R., and Lucarini, G. (2010). Green fluorescent protein as indicator of nonviral transient transfection efficiency in endometrial and testicular biopsies. Microsc Res Tech 73, 229–233. 10.1002/jemt.20779

