## supplementary information for "Integration of Xeno-Free Single-cell Cloning in CRISPR-mediated DNA Editing of Human iPSCs Improves Homogeneity and Methodological Efficiency of Cellular Disease Modelling"

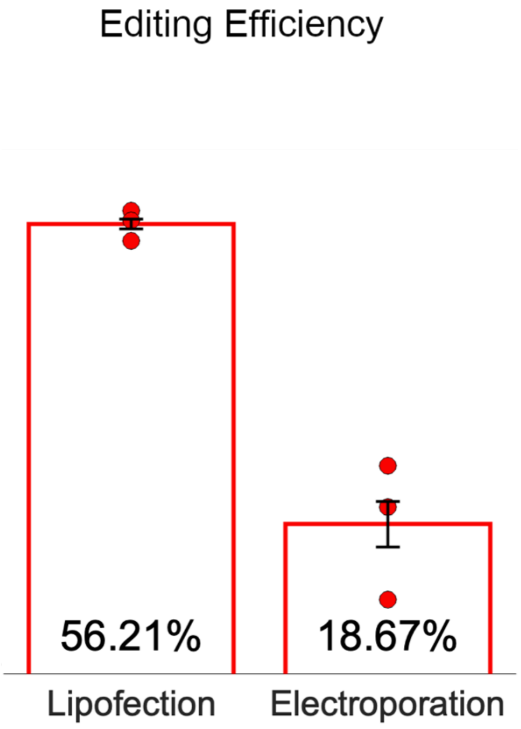


**Supplementary Figure S1 |** Editing efficiency comparison between nucleofection and lipofection

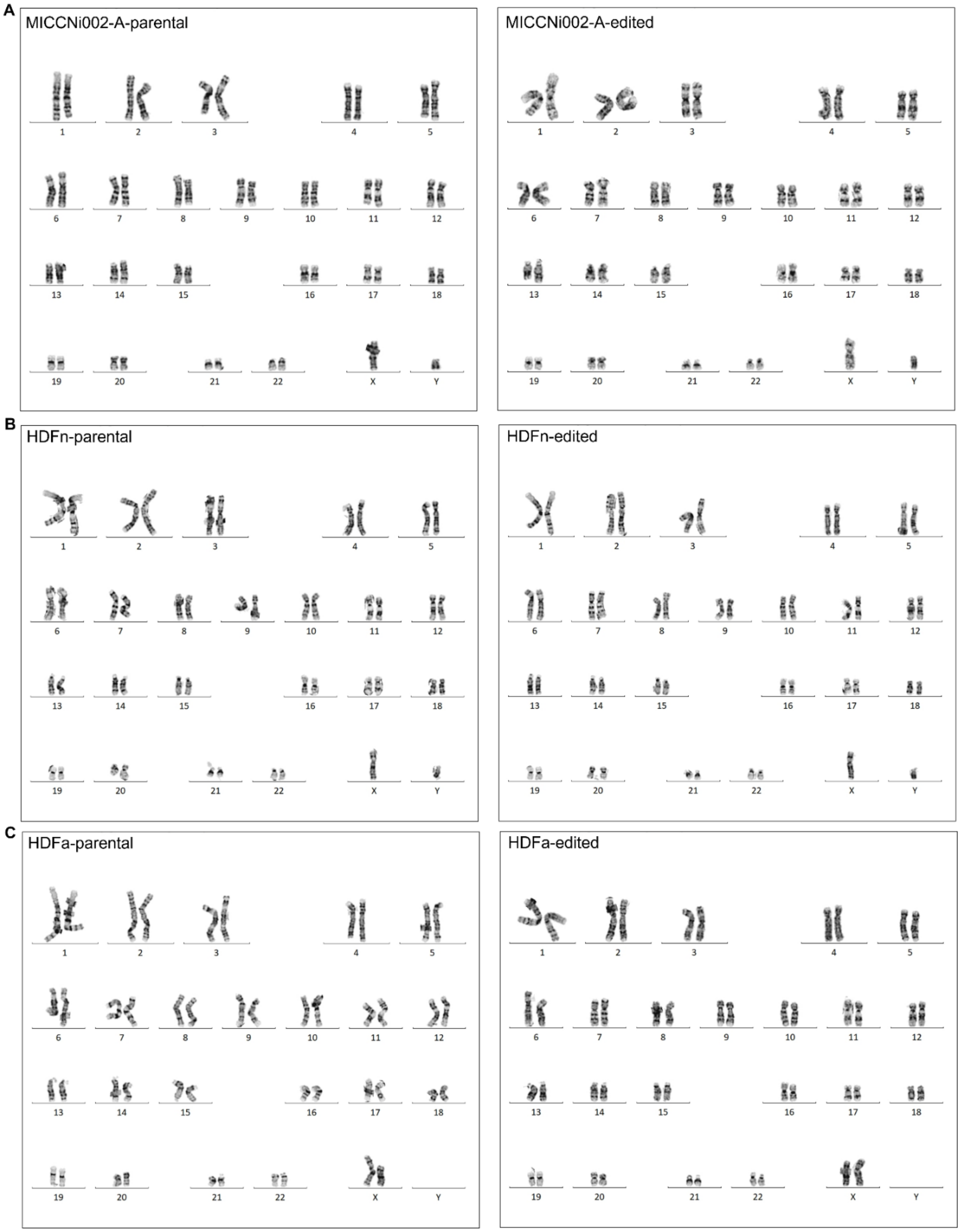


**Supplementary Figure S2 |** Karyotype analysis of edited single-cell clones in MICCNi002-A cell, HDFn and HDFa iPSC lines in comparison to its parental cell line maintaining normal karyotype.

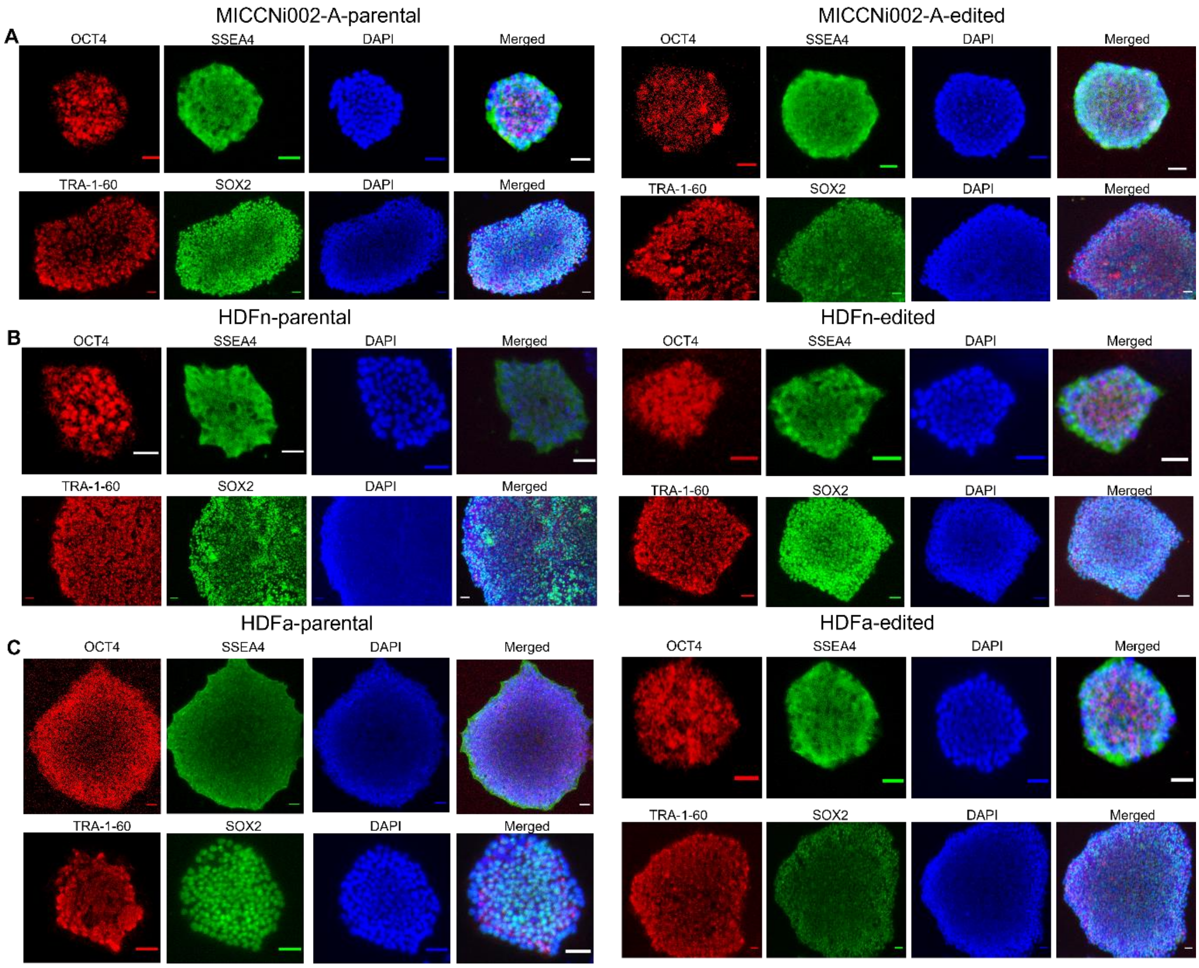


**Supplementary Figure S3** | Assessment of pluripotency of the edited single-cell clones. Pluripotency markers including OCT4, SSEA4, TR-A-1-60 and SOX2 in all three edited iPSC line shows expected expression relative to the parental cell line in (A) MICCi002-A, (B) HDFn and (C) HDFa. Scale bar = 50 µm.

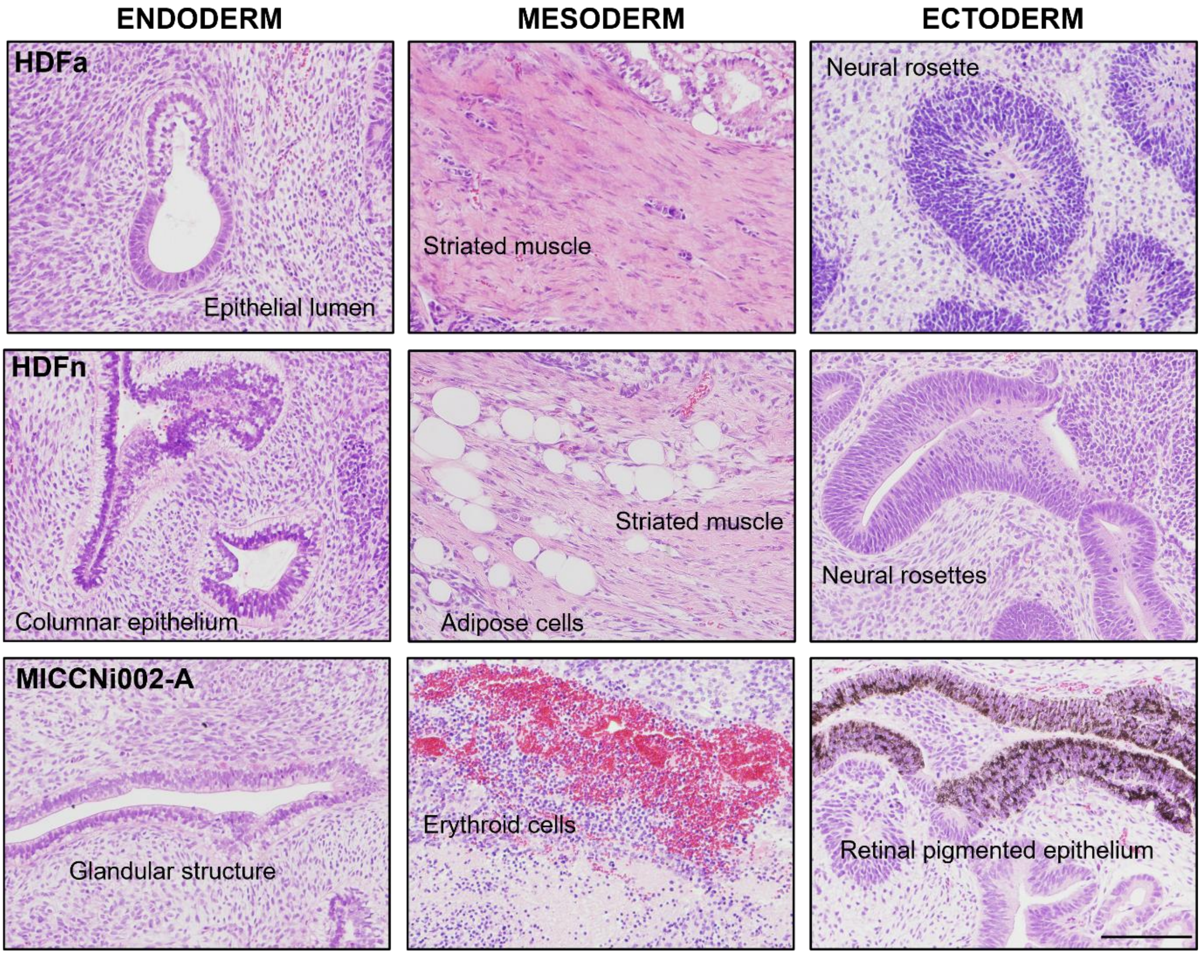


**Supplementary Figure S4 |** Assessment of differentiation potential of edited single-cell clones using teratoma formation assay. Histopathological assessment of the edited single-cell clones of the three iPSC lines is labelled on the left-hand side. Brightfield micrographs of hematoxylin and eosin-stained histological sections show cells that contain derivatives of the three germ layers: endoderm (first column), mesoderm (second column) and ectoderm (third column). The endodermal tissues of the teratomas include epithelial, columnar epithelium, and glandular structure. The mesodermal tissues of the teratomas include striated muscle, adipose cells, and erythroid cells. The ectodermal tissues of the teratomas include neural rosette structures and retinal pigmented epithelium. Scale bar = 200 µm.

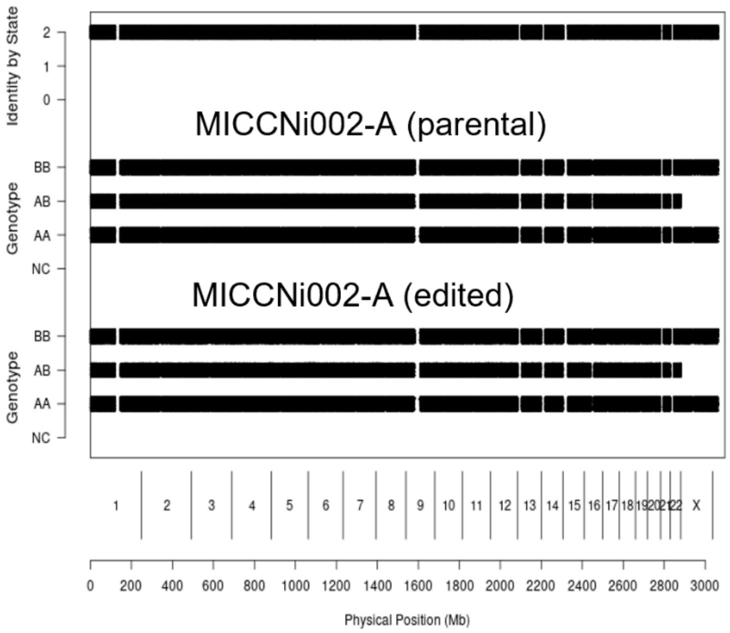
MICCNi002-A (parental)
MICCNi002-A (edited)
Identity by State
Genotype
BB
AB
AA
NC
1 2 3 4 5 6 7 8 9 10 11 12 13 14 15 16 17 18 19 20 21 22 X
0 200 400 600 800 1000 1200 1400 1600 1800 2000 2200 2400 2600 2800 3000
Physical Position (Mb)

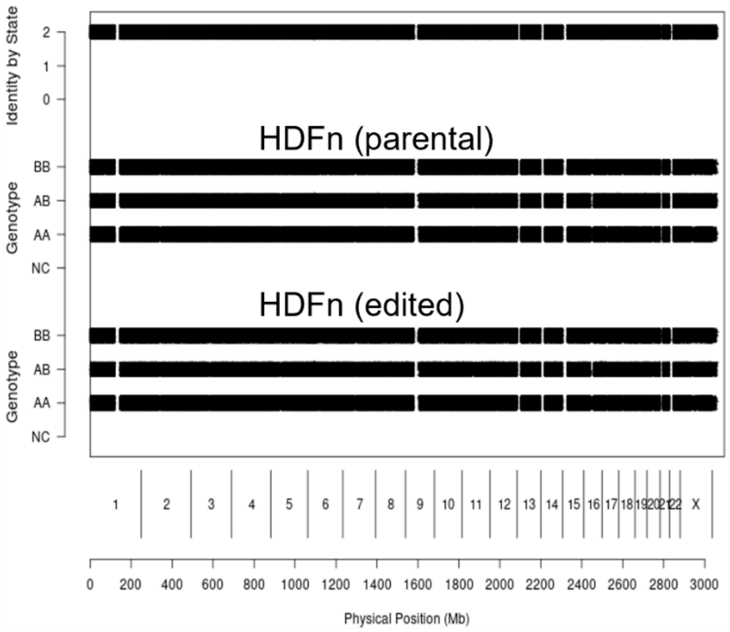
HDFn (parental)
HDFn (edited)
Identity by State
Genotype
BB
AB
AA
NC
1 2 3 4 5 6 7 8 9 10 11 12 13 14 15 16 17 18 19 20 21 22 X
0 200 400 600 800 1000 1200 1400 1600 1800 2000 2200 2400 2600 2800 3000
Physical Position (Mb)

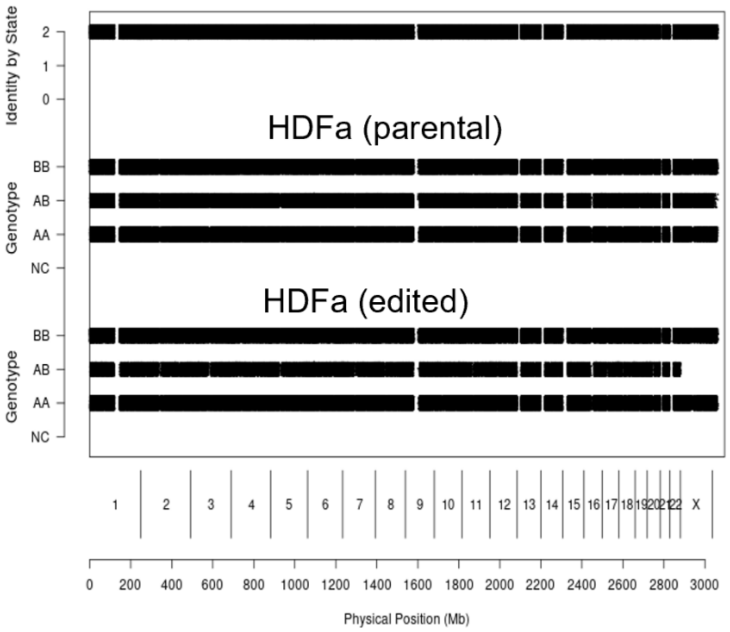
HDFa (parental)
HDFa (edited)
Identity by State
Genotype
BB
AB
AA
NC
1 2 3 4 5 6 7 8 9 10 11 12 13 14 15 16 17 18 19 20 21 22 X
0 200 400 600 800 1000 1200 1400 1600 1800 2000 2200 2400 2600 2800 3000
Physical Position (Mb)

**Supplementary Figure S5 |** Assessment of non-target genomic changes between the edited and parental cell lines using CGH array. SNPduo analysis demonstrated that the edited and parental cell lines are at least 99.9% identical. A comparison of SNP genotypes across the genome is plotted using the SNPduo tool. The top plot shows Identity by State on the y axis with values of 2, 1 and 0, representing two, one and no shared haplotypes, respectively. The middle and lower plots display the B allele frequency on the y axis for each sample.

**Supplementary Table S1 |** Guide RNAs and the donor molecule used in this study.

| Gene | Protospacer Sequence | PAM | Locus position |
|---|---|---|---|
| *FOXP2* | CTAAGCAGCCAATTAGATGC | TGG | chr7: +114426577 |
| *HPRT* | AATTATGGGGATTACTAGGA | AGG | chrX: -134498211 |
| *DUSP6* | CATGCTCCTACCCTATCATT | TGG | chr12: +89348954 |
| *DUSP6* doner | GCCTTACAAGGCCCTAAAGAACACATGCTCCTACCCTATCATTTGA TCCTACCCTATGCGCCTGGAAGTCCTCGAATCCCAGTCCCTG | | |

**Supplementary Table S2 |** Target region primer sequences used in this study.

| Gene | 5' primer for sequencing | 3' primer for sequencing | Fragment length |
|---|---|---|---|
| *FOXP2* | GCATGGTATGTTACTTTGGCTGT | TCCAAAAATCATCCTGTGAGC | 672 bp |
| *HPRT* | AAGAATGTTGTGATAAAAGGTGATGCT | TCAAGGGCATATCCTACAACAA | 698 bp |
| *DUSP6* | CTCTTTGGCTCCTCTATATGCA | AGGTTACCAAGTGCGGAATTG | 300 bp |
| *DUSP6* (HRMA) | CAGCCTTACAAGGCCCTAAA | GGACTGGGATTCGAGGACTT | 87 bp |
